# Induction of neutralizing antibodies against SARS-CoV-2 variants by a multivalent mRNA-lipid nanoparticle vaccine encoding SARS-CoV-2/SARS-CoV Spike protein receptor-binding domains

**DOI:** 10.1101/2022.04.28.489834

**Authors:** Qiong Zhang, Shashi K. Tiwari, Shaobo Wang, Lingling Wang, Wanyu Li, Lingzhi Zhang, Stephen A. Rawlings, Yong Cheng, Jesse V. Jokerst, Tariq M. Rana

## Abstract

To address the need for multivalent vaccines against *Coronaviridae* that can be rapidly developed and manufactured, we compared antibody responses against SARS-CoV, SARS-CoV-2, and several variants of concern in mice immunized with mRNA-lipid nanoparticle vaccines encoding homodimers or heterodimers of SARS-CoV/SARS-CoV-2 receptor-binding domains. All vaccine constructs induced robust anti-viral antibody responses, and the heterodimeric vaccine elicited an IgG response capable of cross-neutralizing SARS-CoV, SARS-CoV-2 Wuhan-Hu-1, B.1.351 (beta), and B.1.617.2 (delta) variants.

## Main text

Beta-coronaviruses (beta-CoV) such as Middle East respiratory syndrome-associated coronavirus (MERS-CoV), severe acute respiratory syndrome coronavirus (SARS-CoV), and SARS-CoV-2, the causative agent of the current COVID-19 pandemic, are associated with high mortality rates worldwide^1,2^. Experience with the Pfizer-BioNTech and Moderna mRNA-based SARS-CoV-2 vaccines approved in 2020/2021 underscores the importance of rapid deployment of vaccines for effective control measures ^3,4^. However, the emergence of more infectious and pathogenic variants of SARS-CoV-2 with enhanced immune escape has highlighted the need for multivalent vaccines that promote immunity to multiple SARS-CoV-2 variants for the current pandemic and, more broadly, to multiple members of the beta-CoV family. In this regard, mRNA vaccines have several advantages over protein-based or inactivated virus-based vaccines, including the feasibility of rapid design and synthesis, low-cost manufacture, and the availability of real-world clinical data supporting the safety of the mRNA platform in humans^5^.

The C-terminal receptor-binding domain (RBD) of the SARS-CoV-2 Spike glycoprotein interacts with angiotensin-converting enzyme 2 (ACE2), the human SARS-CoV-2 receptor, and thus plays a critical role in infection. Both the full-length surface Spike protein and the RBD are potent inducers of neutralizing antibodies and cellular immunity^6^. However, the RBD is also the site of mutations in recently emerged SARS-CoV-2 variants of concern (VOC), including B.1.351 (beta) and B.1.617.2 (delta)^7^. Some of these mutations effectively reduce the neutralizing capacity of antibodies elicited by the current vaccines, resulting in increased transmissibility and/or pathogenicity ^8,9^. Therefore, development of multivalent vaccines must bear in mind the need for broad reactivity to diminish immune escape and protect against rapidly emerging variants^2,4,6,10,13^.

Recent work showed that homodimerization of beta-CoV RBDs increases their stability and immunogenicity^10^, suggesting that homodimers or heterodimers of different beta-CoV RBDs might enhance their ability to elicit cross-reactive humoral and cellular immune responses. Therefore, in the present study, we designed and tested the immunogenicity of four candidate RBD mRNA-lipid nanoparticle (mRNA-LNP) vaccines in mice. We compared the vaccines’ ability to elicit crossreactive antibodies against not only the donor SARS-CoV and SARS-CoV-2 strains but also the recently emerged B.1.351 and B.1.617.2 VOC that exhibit enhanced immune evasiveness.

We designed five mRNA constructs encoding (1) green fluorescent protein (control), (2) SARS-CoV-2 Wuhan-Hu-1 reference strain RBD (R319-K537), (3) SARS-CoV CUHK-W1 RBD (R306–K523), (4) SARS-CoV-2 RBD homodimer, and (5) SARS-CoV/SARS-CoV-2 RBD heterodimer (Fig. 1a). The mRNAs contained an N-terminal human IgE signal peptide to promote secretion; a modified nucleoside N1-methylpseudouridine to increase translation efficiency and reduce activation of the innate immune response^14^; and a cap1 modification at the end of the 5-untranslated region (UTR) and a 3’-UTR poly(A) tail to increase stability and translation efficiency (Supplementary Fig. 1). Transfection of HEK293T cells with the RBD-encoding mRNAs resulted in high expression of each recombinant RBD, as determined by western blot analysis with a cross-reactive anti-SARS-CoV/SARS-CoV-2 RBD antibody (Fig. 1b), which confirmed the competency of the mRNAs to be translated *in vivo.* The RBD mRNAs were then mixed with a combination of lipids optimized to form lipid nanoparticles (LNPs), a commonly used simple and effective delivery vehicle for mRNA vaccines *in vivo*^15^. Measurement of the diameters of multiple batches of RBD mRNA-LNPs by dynamic light scattering (DLS) revealed minimal batch-to batch variation and an average particle diameter of 120 nm (Fig. 1c), a size that results in efficient tissue penetration and cellular uptake ^16^.

**Figure 1.**
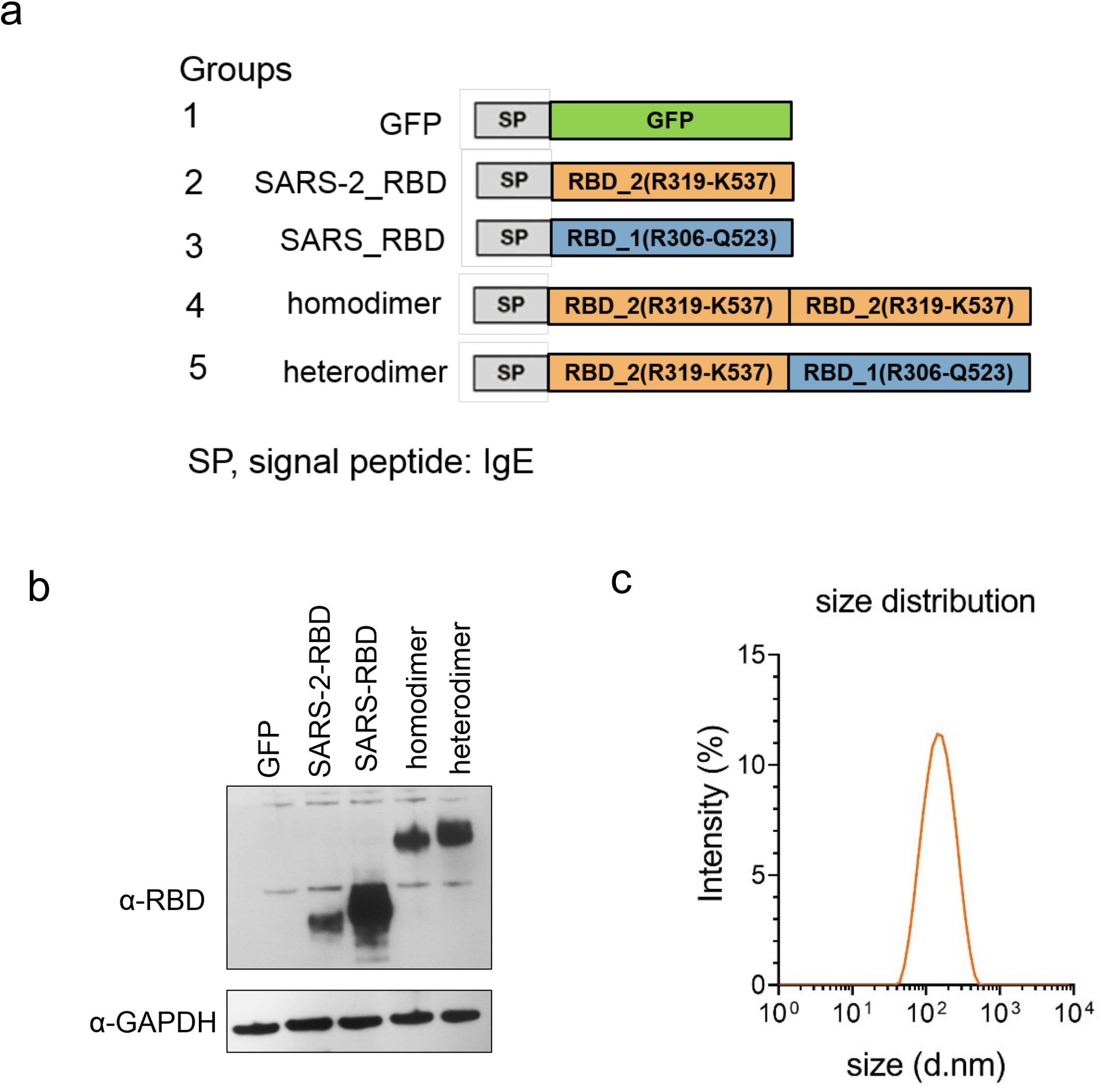
Construction and characterization of multivalent RBD mRNA vaccines. **a** Schematic of the mRNA components of the vaccines. mRNAs encoded the signal peptide of human IgE followed by one or two copies of the receptor-binding domains (RBDs) of SARS-CoV-2 (Wuhan-Hu-1) or SARS-CoV (CUHK-W1) strains. A GFP-expressing vaccine was constructed as a control. **b** Western blot analysis of mRNA-encoded RBD protein expression. Each mRNA was *in vitro* transcribed and transfected into HEK293T cells for 24 h. Brefeldin A (5.0 μg/mL) was added to the cells at 8 h post-transfection to block protein secretion before cell lysis. Blots were probed with a rabbit anti-Spike antibody that recognizes both SARS-CoV and SARS-CoV-2 RBDs. GAPDH was probed as a loading control. **c** Distribution of RBD mRNA-LNP particle diameters measured by dynamic light scattering. For b and c, data from one experiment representative of three independent experiments are shown.

To evaluate the immunogenicity of the RBD mRNA-LNPs, groups of male C57BL/6J mice were immunized intramuscularly with 10 μg of each vaccine in 80 μL phosphate-buffered saline on days 0 and 14 (Fig. 2a). Mice were bled 2 weeks after boosting and sera were analyzed for antibody production by direct binding ELISAs using SARS-CoV-2 and SARS-CoV purified Spike protein-coated plates. Notably, all four RBD mRNA-LNP vaccines elicited high titers of IgG reactive against both Spike proteins (Fig. 2b and c). As expected, the monomer mRNA-LNPs (Groups 2 and 3) showed preferential reactivity against the immunizing Spike protein (Fig. 2b and c); however, the difference in titers was not substantial (mean endpoint titers: Group 2 = 10^4.6^, Group 3 = 10^5.1^). Similarly, the mean anti-Spike protein IgG titers elicited by the SARS-CoV-2 monomer and homodimer mRNA-LNPs were not significantly different (Group 2 = 10^5.1^, Group 4 = 10^5.3^; Fig. 2b), suggesting that dimerization did not increase the immunogenicity of SARS-CoV-2 RBD in the context of these mRNA-LNPs vaccines. Notably, however, the heterodimer mRNA-LNP (Group 5, 10^6.2^) induced a higher titer of anti-SARS-CoV-2 Spike protein IgG compared with the other three vaccines and efficiently elicited antibodies against SARS-CoV Spike protein (Fig. 2b and c). The heterodimeric mRNA-LNP elicited strong and similar IgG2a and IgG1 responses to SARS-CoV-2 Spike protein (Fig. 2d–f), indicative of a balanced immune Th1/Th2 response ^17^.

**Figure 2.**
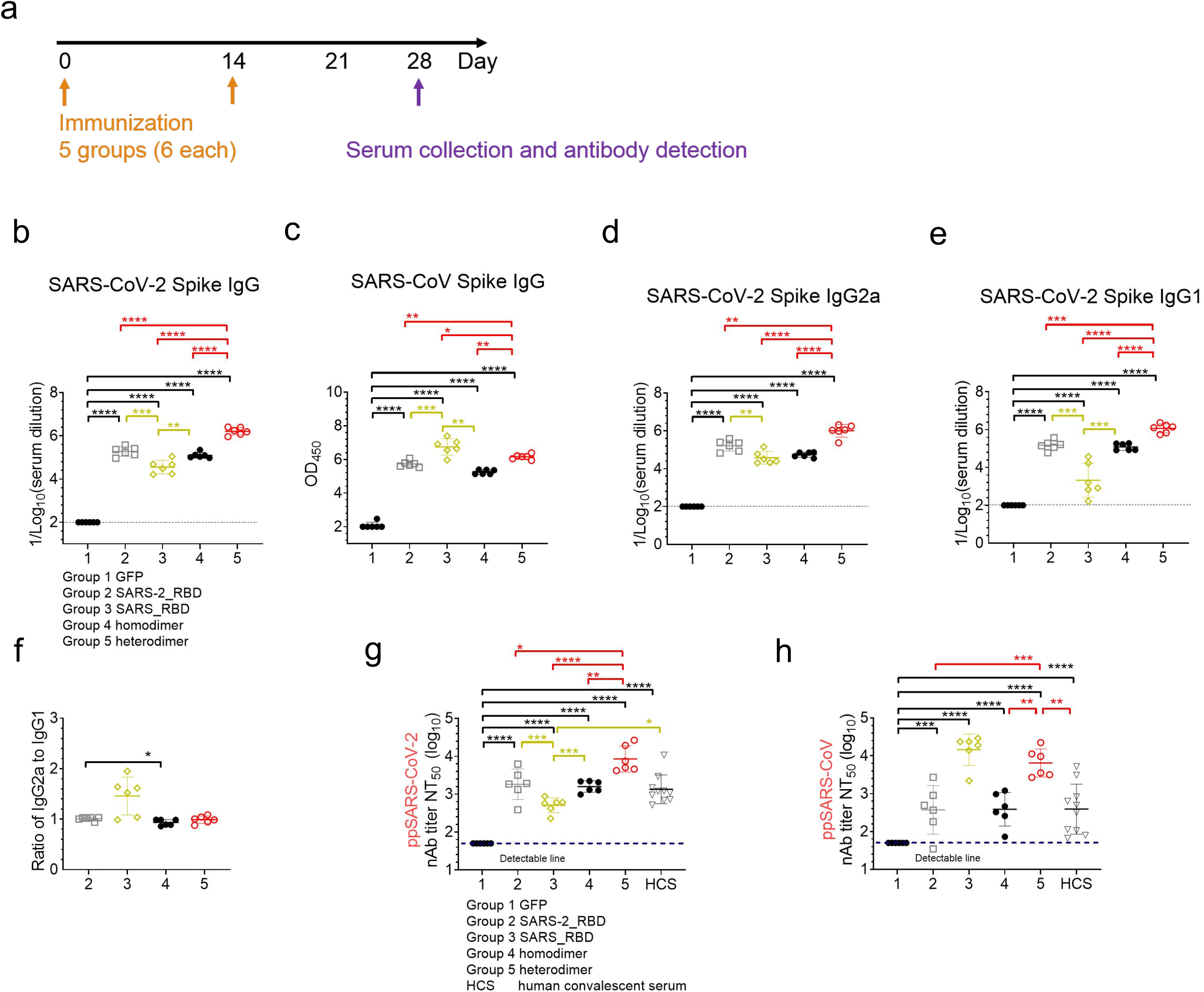
Antibody responses in SARS-CoV and SARS-CoV-2 RBD mRNA-LNP-vaccinated mice. **a** Experimental protocol. Groups of BALB/c mice (n=6) were primed and boosted 2 weeks apart by intramuscular injection of 10 μg of each mRNA-LNP vaccine. Mice were bled on day 28 and sera were prepared. **b–f** Serum titers of SARS-CoV-2 Spike protein-specific (b, d–f) and SARS-CoV Spike protein-specific (c) IgG (b, c), IgG2a (d), and IgG1 (e). The ratio of IgG2a to IgG1 was calculated from the data presented in d and e. **g, h** Serum 50% neutralizing antibody titers (NT_50_) against SARS-CoV-2 (g) and SARS-CoV (h) pseudovirus infection of Vero cells. NT_50_ represents the titer required for 50% inhibition of maximal infection. Dotted lines indicate the limit of detection. Mean ± standard deviation. *\*p* <0.05, *\*\*p* < 0.01, *\*\*\*p* < 0.001, *\*\*\*\*p* < 0.0001, by two-sided *t*-test.

To assess the neutralizing activity of the RBD mRNA-LNP-elicited antibodies, we used a luciferase-based chimeric vesicular stomatitis virus and SARS-CoV pseudovirus (VSVΔG-luc-SARS) assay that quantifies infection enzymatically (Figs. 2g and h). The pattern of neutralizing antibody production in mRNA-LNP-immunized mice was similar to that observed with Spike protein-binding antibodies. Thus, all four RBD mRNA vaccines induced antibodies that effectively neutralized infection of Vero cells with SARS-CoV and SARS-CoV-2-based pseudoviruses (Figs. 2g and h). The NT_50_ (50% neutralization titer) for sera from mice vaccinated with SARS-CoV-2 monomeric, SARS-CoV monomeric, SARS-CoV-2 homodimeric, and SARS-CoV-2 heterodimeric RBD mRNA-LNPs were 1818, 510, 1586, and 8498, respectively, against SARS-CoV-2 pseudovirus (Fig. 2g) and 374, 14510, 385, and 6454, respectively, against the SARS-CoV pseudovirus (Fig. 2h). These data demonstrate that, although the monomeric and heterodimeric SARS-CoV-2 mRNA-LNPs induced comparable Spike protein-binding IgG responses, the heterodimeric vaccine showed a clearly superior ability to induce neutralizing antibodies against both strains.

Antibody responses induced by the two SARS-CoV-2 mRNA vaccines currently in use (mRNA-1273, Moderna(Baden, 2021 #39); BNT162b2, Pfizer-BioNTech^4^) exhibit poorer neutralizing activity against emerging VOCs, including an approximately 30-fold reduction in activity against B.1.351^18^. Therefore, we examined the neutralizing activity of sera from mice immunized with the heterodimeric RBD mRNA-LNP vaccine. We first performed ELISA assays to examine binding and found no difference in antibody binding titers to Wuhan-Hu-1 and B.1.351 Spike proteins (Fig. 3a). Neutralizing activity was then measured against VSVΔG/SARS-CoV-2 pseudoviruses based on Wuhan-Hu-1, B.1.351, B.1.617.2, and Wuhan with N501Y or E484K point mutations. N501Y and E484K are key mutations in the RBD region that confer increased transmissibility^19^. The neutralizing activity of sera from SARS-CoV-2 heterodimeric RBD mRNA-vaccinated mice was reduced approximately 5.4-fold, 2.3-fold, and 14.5-fold against B.1.351, Wuhan-N501Y, and Wuhan-E484K pseudoviruses, respectively, compared with the Wuhan pseudovirus (Fig. 3c and d). Similarly, the heterodimeric RBD mRNA-elicited mouse sera exhibited an 11.7-fold reduction in neutralizing activity against B.1.617.2 (delta) (Fig. 3e). The delta lineage was classified as a VOC in May 2021 due to the increased rate of transmission, reduced effectiveness of monoclonal antibody treatment, and reduced susceptibility to neutralizing antibodies.

**Figure 3.**
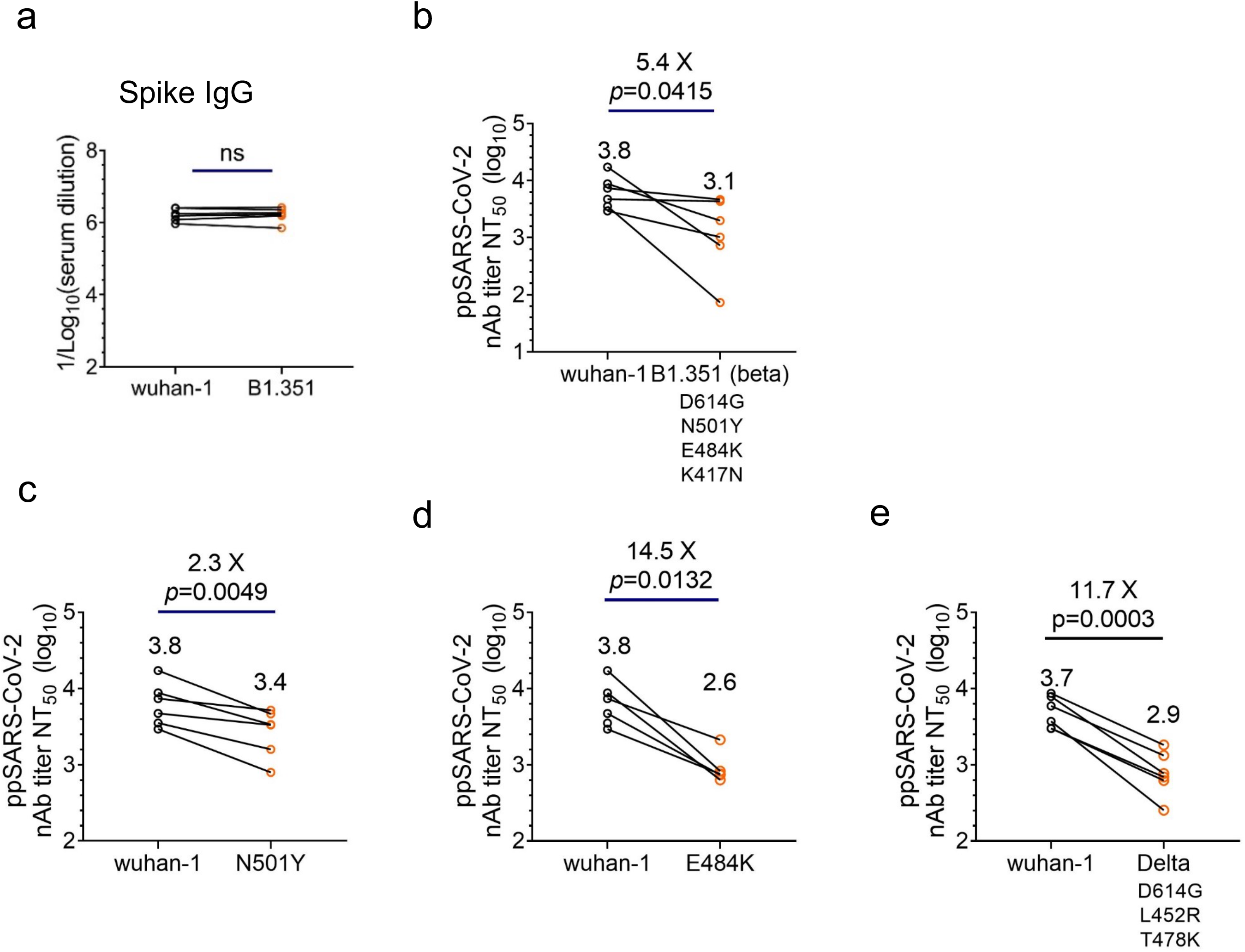
Neutralizing activity against SARS-CoV-2 variants by sera from mice immunized with a heterodimeric RBD mRNA-LNP vaccine. **a** Mice were primed and boosted with the SARS-CoV/SARS-CoV-2 heterodimeric RBD mRNA-LNP as described in Figure 2a and bled 4 weeks after immunization. Direct binding IgG titers were assessed by ELISA against Wuhan-Hu-1 and B.1.351 Spike proteins. **b–e** Neutralizing antibody titers of sera from mice immunized with the RBD heterodimer mRNA-LNP against infection by SARS-CoV-2 Wuhan-Hu-1 compared with (b) B.1.351 (b), Wuhan-N501Y (c), Wuhan-E484K (d), and B.1.617.2 (e) pseudoviruses. Dotted lines indicate the limit of detection. ns, not significant.

Taken together, these data identify a SARS-CoV/SARS-CoV-2 heterodimeric RBD mRNA-LNP vaccine candidate that has the capacity to elicit a strong SARS-CoV/SARS-CoV-2 cross-reactive binding and neutralizing antibody response in mice, including against several SARS-CoV-2 VOCs.

## Methods

### Cell lines

HEK293FT and Vero E6 cells were maintained in Dulbecco’s Modified Eagle’s Medium containing 10% fetal bovine serum (GIBCO). The cell lines were tested and confirmed to be negative for mycoplasma.

### SARS-CoV-2 pseudovirus production

Plasmids encoding the Spike proteins (lacking the C-terminal 19-amino acids) of SARS-CoV-1 (CUHK-W1), SARS-CoV-2 (Wuhan-Hu-1), variant B.1.351, variant B.1.617.2, Wuhan-N501Y, and Wuhan-E484K were transfected into 293T cells with Lipofectamine 3000 (ThermoFisher). After 24 h, the cells were infected for 1 h with VSV-G pseudotyped VSV-dG particles at a multiplicity of infection of 5, washed four times to remove remaining particles, and then incubated in complete medium for 24 h. The supernatants containing the pseudoviral particles were collected, centrifuged to remove cell debris, sterile filtered (Millipore Sigma), and stored in aliquots at −80°C.

### mRNA-LNP generation

mRNA vaccines were designed using the SARS-CoV-2 Wuhan-Hu-1 RBD protein sequence (GenBank: MN908947.3) and SARS-CoV CUHK-W1 RBD protein sequence (GenBank: AY278554.2). Five test vaccines were constructed (Fig. 1a) consisting of a C-terminal IgE signal peptide (SP) followed by coding sequences for GFP (control), SARS-CoV-2 RBD (R319-K537), SARS-CoV RBD (R306-Q523), homodimer of SARS-CoV-2 RBD, and heterodimer of SARS-CoV-2 RBD.

The sequences were codon-optimized and cloned into pbluscript, an mRNA production plasmid generated in our lab. As shown in Supplementary Figure and Supplementary Data, the sequences comprised the T7 promoter, 5’ and 3’ untranslated regions of human hemoglobin subunit alpha 1, IgE signal peptide, and the respective RBD coding sequences. The DNA vectors were linearized and the mRNA was synthesized *in vitro* using T7 polymerase (Cellscript, #C-ASF3507), with UTP substituted by m1Ψ-5’-triphosphate (TriLink, #N-1081). A donor methyl group S-adenosylmethionine was added to the methylated capped RNA (cap 0), resulting in a cap 1 structure to increase mRNA translation efficiency (Cellscript, #C-SCCS1710). The poly(A) tail was added using a Poly(A) Tailing Kit (Thermo Fisher Scientific, #74225Z25KU). The mRNA was purified using LiCl (Sigma, SLCC8730, 2 M final concentration). To generate the LNPs, (6Z,9Z,28Z,31Z)-heptatriaconta-6,9,28,31-tetraen-19-yl 4-(dimethylamino)butanoate (D-Lin-MC3-DMA), 1, 2-distearoyl-sn-glycero-3-phosphocholine, cholesterol, and 1,2-dimyristoyl-rac-glycero-3-methoxypolyethylene glycol-2000 (DMG-PEG2000) were combined in ethanol at a molar ratio of 50:10:38.5:1.5. LNPs were formed by a self-assembly process in which the lipid mixture was rapidly mixed with the relevant indicated mRNA in 100 mM sodium acetate (pH 4) and incubated at 37°C for 15 min. The mRNA-LNP was then diluted in PBS to give a final mRNA concentration of 167 μg/mL. To measure mRNA-LNP size, the solution was diluted 10-fold in PBS and analyzed using a dynamic light scattering machine (Malvern NANO-ZS90 Zetasizer). The experiments were performed with three batches of each mRNA-LNP vaccine, all of which were comparable in size.

### Verification of protein-coding capability of vaccine mRNAs

To confirm that the synthesized mRNAs could be translated into GFP or RBD proteins, 293FT cells were seeded at 3 × 10^5^ cells/mL in 6-well plates, grown for 24 h, and then transfected with 1 mg mRNA per well using Lipofectamine 3000 (Invitrogen) according to the manufacturer’s instructions. To prevent secretion of proteins, cells were incubated with the protein transport inhibitor brefeldin A (5.0 μg/mL) for 8 h after transfection. The transfected cells were cultured at 37°C for 24 h and then collected and lysed using protein lysis buffer (Thermo Fisher Scientific, Cat #: 87787). Aliquots of lysate (20 μg protein) were resolved on 4–12% NuPAGE precast gels (Thermo Fisher Scientific) and transferred to PVDF membranes. RBD protein expression was analyzed using a rabbit polyclonal antibody SARS-CoV-2 Spike RBD Antibody (HRP) (Sino Biological, #40592-T62), which cross-reacts with SARS-CoV RBD protein. Glyceraldehyde 3-phosphate dehydrogenase (GAPDH) was probed as a loading control. At least three batches of mRNA were used in these experiments, and no significant variability in protein expression was noted.

### Immunization of mice with RBD mRNA-LNPs

Male C57BL/6J mice (aged 4–5 weeks) were purchased from the Jackson Laboratory and housed according to the regulatory standards of the University of California, San Diego. Mice were randomly allocated to experimental groups. The mice were primed by intramuscular injection of mRNA-LNP (10 μg mRNA in 80 μL PBS) into the quadriceps muscle and then boosted 2 weeks later with the same mRNA-LNP dose and administration route. Blood was collected *via* cardiac puncture 2 weeks after the boost (4 weeks post-immunization) and serum was prepared and stored at −80°C until analyzed.

### SARS-CoV Spike protein-specific ELISAs

ELISAs were designed to quantify SARS-CoV-2 Spike protein-reactive total IgG, IgG2a, and IgG1, as well as SARS-CoV Spike protein-reactive total IgG. Recombinant SARS-CoV-2, SARS-CoV, or B.1.351 Spike proteins (Sino Biological) were diluted to 200 ng/mL in 50 mM sodium carbonate buffer (pH 9.6), added to 96-well EIA/RIA plates (Corning) at 100 μL/well, and incubated overnight at 4°C. The plates were washed with PBS containing 0.5% Tween-20 (PBST) and blocked with 5% bovine serum albumin (BSA) in PBS for 30 min at 37°C. Mouse serum samples were serially diluted 5-fold in PBST containing 1% BSA and 100 μL was added to each well and incubated for 2 h at 37°C. The plates were washed three times with PBST and incubated with horseradish peroxidase-conjugated goat anti-mouse IgG (Abcam, ab6789, 1:10,000), goat anti-mouse IgG1 (Abcam, ab97240, 1:10,000), or goat anti-mouse IgG2a (Abcam, ab97245, 1:10,000) for 1 h. The plates were washed with PBST, color was developed by addition of 3’,5,5’-tetramethylbenzidine (TMB) substrate, and the reaction was stopped by addition of 2 M HCl. The absorbance (optical density, OD) at 450 nm was measured using a microplate reader (BioTeK). The endpoint dilution titer was defined as the highest serum dilution giving an OD >2-fold the background OD of control wells consisting of diluted serum.

### Neutralizing antibody assay with pseudoviruses

The neutralizing antibody titer of serum samples from vaccinated mice was determined by measuring the ability to block infection of Vero cells by chimeric VSVΔG-luc-SARS pseudoviruses (SARS-CoV, SARS-CoV-2 Wuhan-Hu-1, B.1.351, B.1.617.2, Wuhan-N501Y, or Wuhan-E484K). Vero cells were seeded at 2 × 10^5^ cells/mL of 100 μL/well in 96-well plates and cultured overnight at 37°C. Serum samples were heat inactivated at 56°C for 30 min, serially diluted 3-fold in DMEM medium, mixed with VSVΔG-luc-SARS at 100 TCID50 (50% tissue culture infectious dose), and incubated at 37°C for 1 h. The mixture was then added to the plated Vero cells at 100 μL/well and incubated for 24 h at 37°C. All samples were assayed in triplicate and controls consisting of Vero cells cultured alone or with pseudovirus without serum preincubation were tested in parallel. After 24 h, Vero cells were lysed and luciferase activity was measured using a Bright-Glo firefly luciferase kit (Promega). The 50% neutralization titer (NT_50_) was calculated as the reciprocal of the highest serum dilution that gave a 50% reduction in luciferase signal compared with the negative control samples. NT_50_ values were calculated using GraphPad Prism 8.0.

### Statistical analysis

Data are presented as the mean and standard deviation, and symbols represent individual samples or mice. Group means were compared using an unpaired *t*-test (Fig. 2), or a nonparametric two-tailed Student’s *t*-test (Fig. 3). Statistical analyses were conducted using GraphPad Prism 8.0. A *P* value <0.05 was considered to be statistically significant.

## Acknowledgements

We thank members of the Rana lab for helpful assistance. JVJ acknowledges NIH support under R21 AI157957. This work was supported in part by NIH grants (AI125103, DA046171 and DA039562).

## Author contributions

QZ designed and performed the experiments, analyzed the data, and wrote the manuscript; ST, designed and performed the experiments, analyzed the data; LW, SW, WL, LZ performed experiments and analyzed the data; SAR provided reagents, YC and JVJ assisted in LNP measurements; TMR conceived the overall project and contributed to the experimental design, data analysis, data interpretation, and manuscript writing.

## Data availability

The datasets that support the findings of this study are available from the corresponding author upon reasonable request.

## Supplementary information

Supplementary Data. DNA sequences for *in vitro* transcription of vaccine mRNAs

**Supplementary Figure 1.**
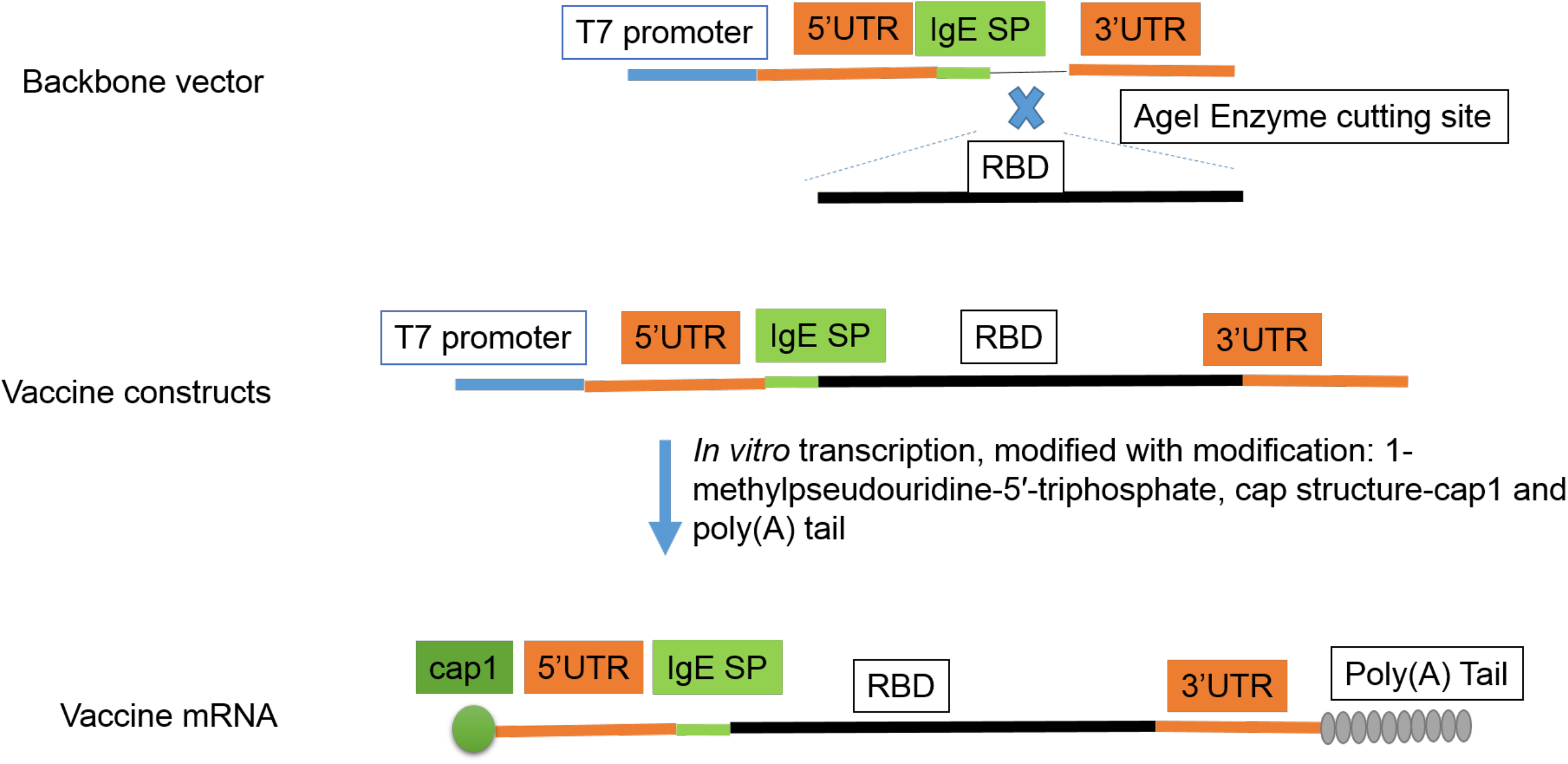
Design of mRNA constructs.

## References

1 Cui, J., Li, F. & Shi, Z. L. Origin and evolution of pathogenic coronaviruses. Nat Rev Microbiol 17, 181–192, doi:10.1038/s41579-018-0118-9 (2019).

2 Hu, B., Guo, H., Zhou, P. & Shi, Z. L. Characteristics of SARS-CoV-2 and COVID-19. Nat Rev Microbiol 19, 141–154, doi:10.1038/s41579-020-00459-7 (2021).

3 Baden, L. R. et al. Efficacy and Safety of the mRNA-1273 SARS-CoV-2 Vaccine. The New England journal of medicine 384, 403–416, doi:10.1056/NEJMoa2035389 (2021).

4 Polack, F. P. et al. Safety and Efficacy of the BNT162b2 mRNA Covid-19 Vaccine. The New England journal of medicine 383, 2603–2615, doi:10.1056/NEJMoa2034577 (2020).

5 Chaudhary, N., Weissman, D. & Whitehead, K. A. mRNA vaccines for infectious diseases: principles, delivery and clinical translation. Nature reviews. Drug discovery, doi:10.1038/s41573-021-00283-5 (2021).

6 Kyriakidis, N. C., Lopez-Cortes, A., Gonzalez, E. V., Grimaldos, A. B. & Prado, E. O. SARS-CoV-2 vaccines strategies: a comprehensive review of phase 3 candidates. NPJ vaccines 6, 28, doi:10.1038/s41541-021-00292-w (2021).

7 Zhou, W. & Wang, W. Fast-spreading SARS-CoV-2 variants: challenges to and new design strategies of COVID-19 vaccines. Signal transduction and targeted therapy 6, 226, doi:10.1038/s41392-021-00644-x (2021).

8 Harvey, W. T. et al. SARS-CoV-2 variants, spike mutations and immune escape. Nat Rev Microbiol 19, 409–424, doi:10.1038/s41579-021-00573-0 (2021).

9 Lopez Bernal, J. et al. Effectiveness of Covid-19 Vaccines against the B.1.617.2 (Delta) Variant. The New England journal of medicine 385, 585–594, doi:10.1056/NEJMoa2108891 (2021).

10 Dai, L. et al. A Universal Design of Betacoronavirus Vaccines against COVID-19, MERS, and SARS. Cell 182, 722–733 e711, doi:10.1016/j.cell.2020.06.035 (2020).

11 Zhang, N. N. et al. A Thermostable mRNA Vaccine against COVID-19. Cell 182, 1271–1283 e1216, doi:10.1016/j.cell.2020.07.024 (2020).

12 Huang, Q. et al. A single-dose mRNA vaccine provides a long-term protection for hACE2 transgenic mice from SARS-CoV-2. Nature communications 12, 776, doi:10.1038/s41467-021-21037-2 (2021).

13 Laczko, D. et al. A Single Immunization with Nucleoside-Modified mRNA Vaccines Elicits Strong Cellular and Humoral Immune Responses against SARS-CoV-2 in Mice. Immunity 53, 724–732 e727, doi:10.1016/j.immuni.2020.07.019 (2020).

14 Kariko, K., Buckstein, M., Ni, H. & Weissman, D. Suppression of RNA recognition by Toll-like receptors: the impact of nucleoside modification and the evolutionary origin of RNA. Immunity 23, 165–175, doi:10.1016/j.immuni.2005.06.008 (2005).

15 Ickenstein, L. M. & Garidel, P. Lipid-based nanoparticle formulations for small molecules and RNA drugs. Expert opinion on drug delivery 16, 1205–1226, doi:10.1080/17425247.2019.1669558 (2019).

16 Foroozandeh, P. & Aziz, A. A. Insight into Cellular Uptake and Intracellular Trafficking of Nanoparticles. Nanoscale Res Lett 13, doi:Artn33910.1186/S11671-018-2728-6 (2018).

17 Stevens, T. L. et al. Regulation of antibody isotype secretion by subsets of antigen-specific helper T cells. Nature 334, 255–258, doi:10.1038/334255a0 (1988).

18 Garcia-Beltran, W. F. et al. Multiple SARS-CoV-2 variants escape neutralization by vaccine-induced humoral immunity. Cell 184, 2523, doi:10.1016/j.cell.2021.04.006 (2021).

19 Zhou, D. et al. Evidence of escape of SARS-CoV-2 variant B.1.351 from natural and vaccine-induced sera. Cell 184, 2348–2361 e2346, doi:10.1016/j.cell.2021.02.037 (2021).

